# A natural mutation between SARS-CoV-2 and SARS-CoV determines neutralization by a cross-reactive antibody

**DOI:** 10.1101/2020.09.21.305441

**Authors:** Nicholas C. Wu, Meng Yuan, Sandhya Bangaru, Deli Huang, Xueyong Zhu, Chang-Chun D. Lee, Hannah L. Turner, Linghang Peng, Linlin Yang, David Nemazee, Andrew B. Ward, Ian A. Wilson

**Affiliations:** Department of Biochemistry, University of Illinois at Urbana-Champaign, Urbana, IL 61801, USA; Carl R. Woese Institute for Genomic Biology, University of Illinois at Urbana-Champaign, Urbana, IL 61801, USA; Department of Integrative Structural and Computational Biology, The Scripps Research Institute, La Jolla, CA 92037, USA; Department of Immunology and Microbiology, The Scripps Research Institute, La Jolla, CA 92037, USA; The Skaggs Institute for Chemical Biology, The Scripps Research Institute, La Jolla, CA, 92037, USA

## Abstract

Epitopes that are conserved among SARS-like coronaviruses are attractive targets for design of cross-reactive vaccines and therapeutics. CR3022 is a SARS-CoV neutralizing antibody to a highly conserved epitope on the receptor binding domain (RBD) on the spike protein that can cross-react with SARS-CoV-2, but with lower affinity. Using x-ray crystallography, mutagenesis, and binding experiments, we illustrate that of four amino acid differences in the CR3022 epitope between SARS-CoV-2 and SARS-CoV, a single mutation P384A fully determines the affinity difference. CR3022 does not neutralize SARS-CoV-2, but the increased affinity to SARS-CoV-2 P384A mutant now enables neutralization with a similar potency to SARS-CoV. We further investigated CR3022 interaction with the SARS-CoV spike protein by negative-stain EM and cryo-EM. Three CR3022 Fabs bind per trimer with the RBD observed in different up-conformations due to considerable flexibility of the RBD. In one of these conformations, quaternary interactions are made by CR3022 to the N-terminal domain (NTD) of an adjacent subunit. Overall, this study provides insights into antigenic variation and potential for cross-neutralizing epitopes on SARS-like viruses.

## INTRODUCTION

The ongoing COVID-19 pandemic, which is caused by the new coronavirus SARS-CoV-2, continues to escalate. Investigating the immunogenicity and antigenicity of SARS-CoV-2 is of great importance for vaccine and therapeutic development. The major antigen of coronavirus is the spike (S) glycoprotein, which is expressed as a homotrimer on the virus surface. Since the S protein is essential for virus entry through engaging the host receptor and mediating virus-host membrane fusion, many antibodies to the S protein are neutralizing [1-12]. The S proteins of SARS-CoV-2 and SARS-CoV, which caused a global outbreak in 2003, have an amino-acid sequence identity of around 77% [13] that leads to differences in antigenicity in serology studies [14, 15]. Although a few monoclonal antibodies have been discovered that can cross-neutralize SARS-CoV and SARS-CoV-2 [6, 7, 16, 17], they seem to be relatively rare in COVID-19 patients [1, 3, 4, 14]. Thus, the molecular determinants that define the antigenic differences and similarities between SARS-CoV-2 and SARS-CoV need further exploration.

CR3022 was previously isolated from a SARS survivor and neutralizes SARS-CoV [18], CR3022 was recently found to also be a cross-reactive antibody that can bind to both SARS-CoV-2 and SARS-CoV [19]. Our recent crystal structure demonstrated that CR3022 targets a highly conserved cryptic epitope on the receptor binding domain (RBD) of the S protein [20]. The CR3022 epitope is exposed only when the RBD is in the “up” but not the “down” conformation on the S protein [20]. A few SARS-CoV-2 antibodies from COVID-19 patients have also recently been shown to target the CR3022 epitope [12, 17, 21], suggesting that it is an important site of vulnerability for the antibody response in SARS-CoV-2 infection. Out of 28 residues in the CR3022 epitope, 24 are conserved between SARS-CoV-2 and SARS-CoV, explaining the cross-reactive binding of CR3022. However, CR3022 has a higher affinity to SARS-CoV than to SARS-CoV-2 (>100-fold difference), and can neutralize SARS-CoV, but not SARS-CoV-2, in a live virus neutralization assay [20]. Therefore, CR3022 provides a good case study to probe antigenic variation between SARS-CoV-2 and SARS-CoV.

We therefore aimed to dissect the molecular basis underlying the difference in binding affinity and neutralization potency of CR3022 to SARS-CoV-2 and SARS-CoV. The crystal structure of SARS-CoV RBD in complex with CR3022 was determined to compare with the corresponding SARS-CoV-2 RBD structure [20]. In combination of mutagenesis and binding experiments, we demonstrate that a single amino-acid difference at residue 384 (SARS-CoV-2 numbering) between the RBDs of SARS-CoV-2 and SARS-CoV can fully explain the affinity difference with CR3022. Moreover, CR3022 is now able to neutralize SARS-CoV-2 P384A with a similar potency to SARS-CoV. We further investigated the molecular recognition of CR3022 to the SARS-CoV-2 spike protein by electron microscopy and found that rotational flexibility of the RBD resulted in antibody binding to different variants of up-conformations of the RBD relative to the spike trimer. Our findings validate the CR3022 epitope as an important site of vulnerability for a cross-neutralizing antibody response. Throughout this study, residues on RBD are numbered according to SARS-CoV-2 numbering unless otherwise stated.

## RESULTS

### P384A increases binding affinity of SARS-CoV-2 RBD to CR3022

The epitope of CR3022 in SARS-CoV-2 and SARS-CoV differs by four residues. We aimed to determine whether amino-acid variants in these four non-conserved residues influence the binding affinity of CR3022 to RBD. Four SARS-CoV-2 RBD mutants, namely A372T, P384A, T430M, and H519N (SARS-CoV-2 numbering), were recombinantly expressed and examined (Figure 1A). These mutants converted the amino-acid sequence of the CR3022 epitope in the SARS-CoV-2 RBD to that of SARS-CoV at each of the four non-conserved residues. While binding of CR3022 mutants A372T (K_D_ = 66 nM), T430M (K_D_ = 64 nM), and H519N (K_D_ = 60 nM) was comparable to wild type (WT) SARS-CoV-2 RBD (K_D_ = 68 nM), binding of CR3022 to the P384A mutant (K_D_ = 1.4 nM) was greatly increased (Figure 1B), akin now to that with the SARS-CoV RBD (K_D_ = 1.0 nM) [20]. Thus, the difference in binding affinity of CR3022 to SARS-CoV-2 RBD versus SARS-CoV RBD can be attributed due to a single amino-acid difference at residue 384.

**Figure 1.**
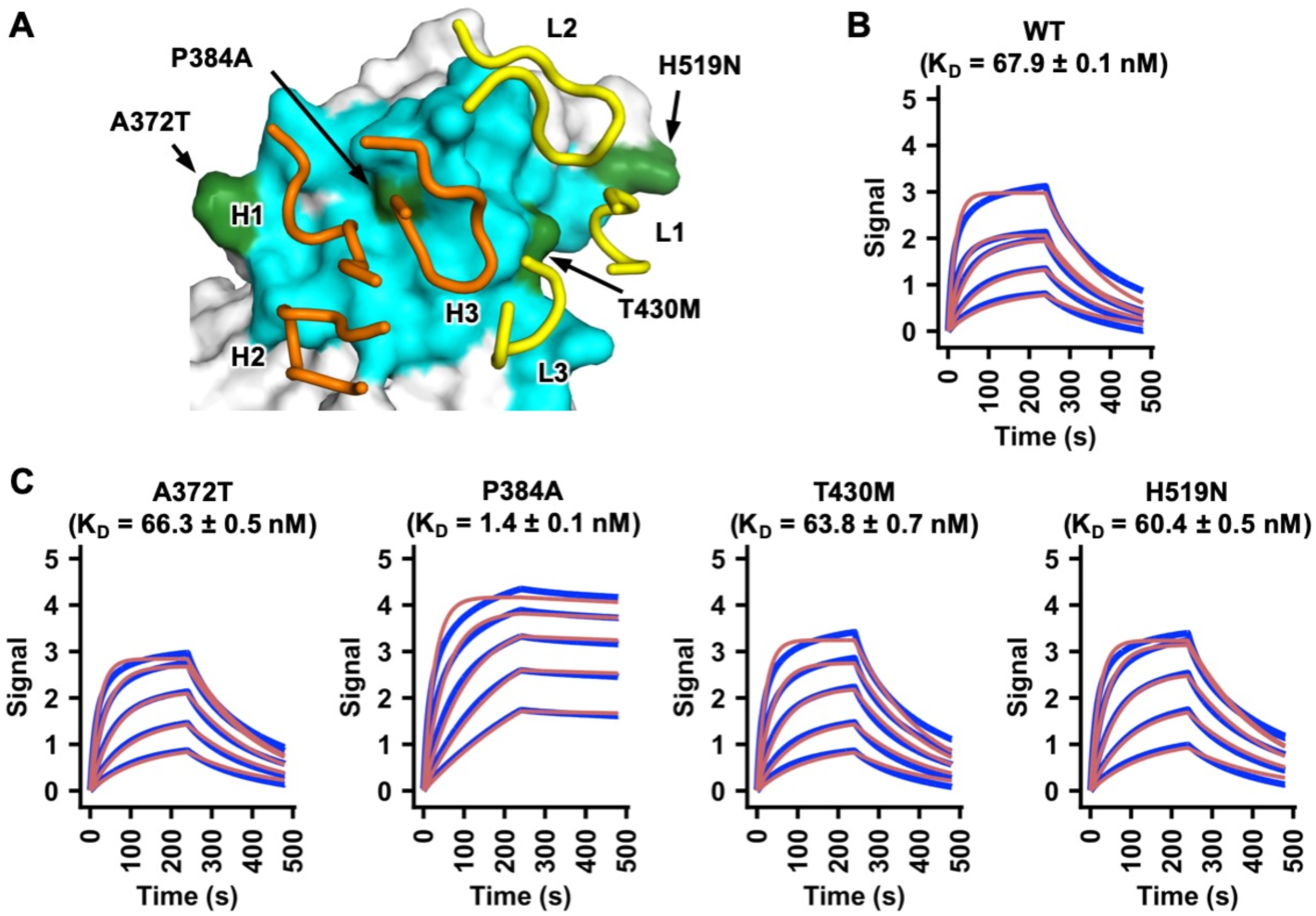
A single P384A substitution increases CR3022 affinity to the SARS-CoV-2 RBD. **(A)** Epitope residues on SARS-CoV RBD are colored in cyan and green. The CR3022 CDR loops that contact the RBD are shown and labeled. Cyan: epitope residues that are conserved between SARS-CoV-2 and SARS-CoV. Green: epitope residues that are not conserved between SARS-CoV-2 and SARS-CoV. Orange: CR3022 heavy chain. Yellow: CR3022 light chain. **(B-C)** Binding of CR3022 Fab to **(B)** wild-type (WT) SARS-CoV-2 RBD and **(C)** different mutants was measured by biolayer interferometry with RBD loaded on the biosensor and Fab in solution. Y-axis represents the response. Dissociation constants (K_D_) for the Fab were obtained using a 1:1 binding model, respectively, which is represented by the red curves. Representative results of two replicates for each experiment are shown. Of note, mammalian cell-expressed RBD was used in the binding experiments in this study, whereas insect cell-expressed RBD was used in our previous study [20]. This difference may explain the slight discrepancy in the K_D_ of CR3022 Fab to SARS-CoV-2 RBD WT between this study and our previous study [20].

### CR3022 neutralizes SARS-CoV-2 P384A but not WT

While CR3022 can neutralize SARS-CoV [18, 20], multiple groups have shown that it does not neutralize SARS-CoV-2 [3, 5, 20, 22]. One possibility is that the affinity of CR3022 to SARS-CoV-2 RBD is not sufficient to confer neutralizing activity. To test this hypothesis, we compared neutralization of SARS-CoV-2 WT and the P384A mutant by CR3022. Consistent with previous studies [3, 5, 20, 22], CR3022 failed to neutralize SARS-CoV-2 WT (Figure 2). However, CR3022 is now able to neutralize the SARS-CoV-2 P384A mutant at an IC_50_ of 3.2 μg/ml, which is comparable to its neutralizing activity to SARS-CoV (IC_50_ of 5.2 μg/ml). This finding validates the CR3022 epitope as a neutralizing epitope in both SARS-CoV-2 and SARS-CoV, provided that the antibody affinity can surpass a threshold for detection of neutralization.

**Figure 2.**
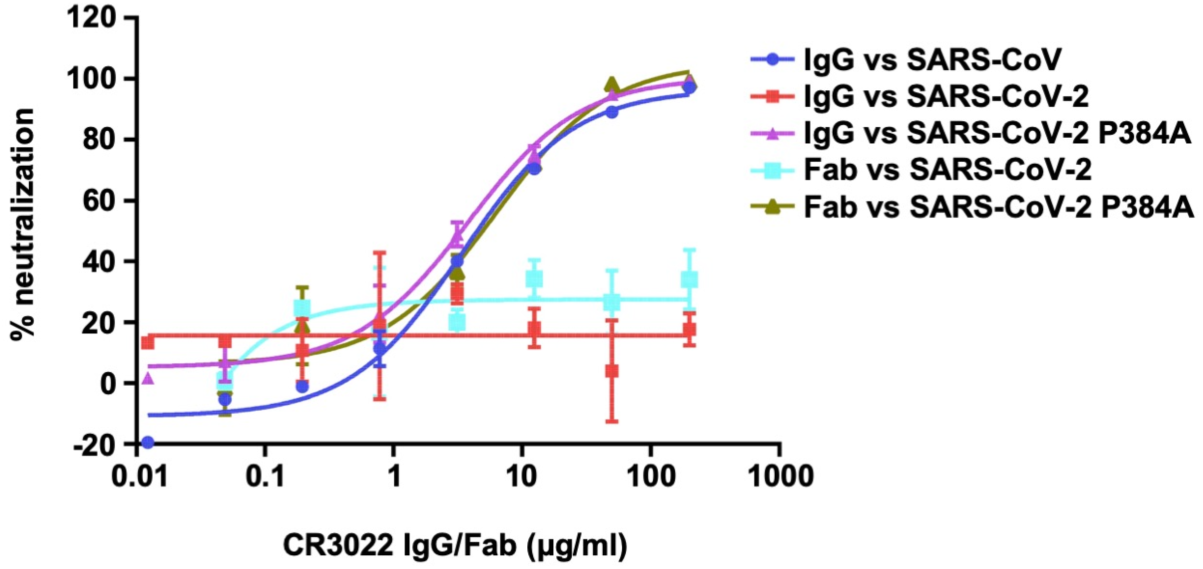
Pseudovirus neutralization assay. The neutralizing activity of CR3022 IgG or Fab to SARS-CoV, SARS-CoV-2, and SARS-CoV-2 P384A mutant was measured in a pseudovirus neutralization assay.

Previous studies have indicated IgG bivalent binding can play an important role in mediating neutralization of SARS-CoV-2, since the neutralization potency for many antibodies is much greater when expressed as IgG rather than Fab [21, 23]. Subsequently, we also tested the neutralizing activity of CR3022 Fab. Interestingly, the CR3022 Fab neutralized SARS-CoV-2 P384A mutant with an IC_50_ of 4.4 μg/ml, which is similar to that of CR3022 IgG (3.2 μg/ml) (Figure 2). This result indicates that CR3022, unlike many other SARS-CoV-2 antibodies [21, 23], does not act bivalently with the S proteins on the virus surface and, hence, neutralization is more sensitive to Fab binding affinity.

### Sequence conservation of residue 384

We then examined the sequence conservation of residue 384 in other SARS-related coronaviruses (SARSr-CoV) strains. Most SARSr-CoV strains have Pro at residue 384, as in SARS-CoV-2. Only those strains that are phylogenetically very close to SARS-CoV, such as bat SARSr-CoV WIV1 and bat SARSr-CoV WIV16, have Ala at residue 384 (Figure 3A). Phylogenetic analysis implies that P384A emerged during the evolution of SARSr-CoV in bats (Figure 3A), which is the natural reservoir of SARSr-CoV [24]. However, it is unclear whether the emergence of P384A is due to neutral drift or positive selection in bats or other species. In addition, given that residue 384 is proximal to the S2 domain when the RBD is in the “down” conformation (Figure 3B), whether P384A can modulate the conformational dynamics of the “up and down” configurations of the RBD in the S trimer and influence the viral replication fitness will require additional studies.

**Figure 3.**
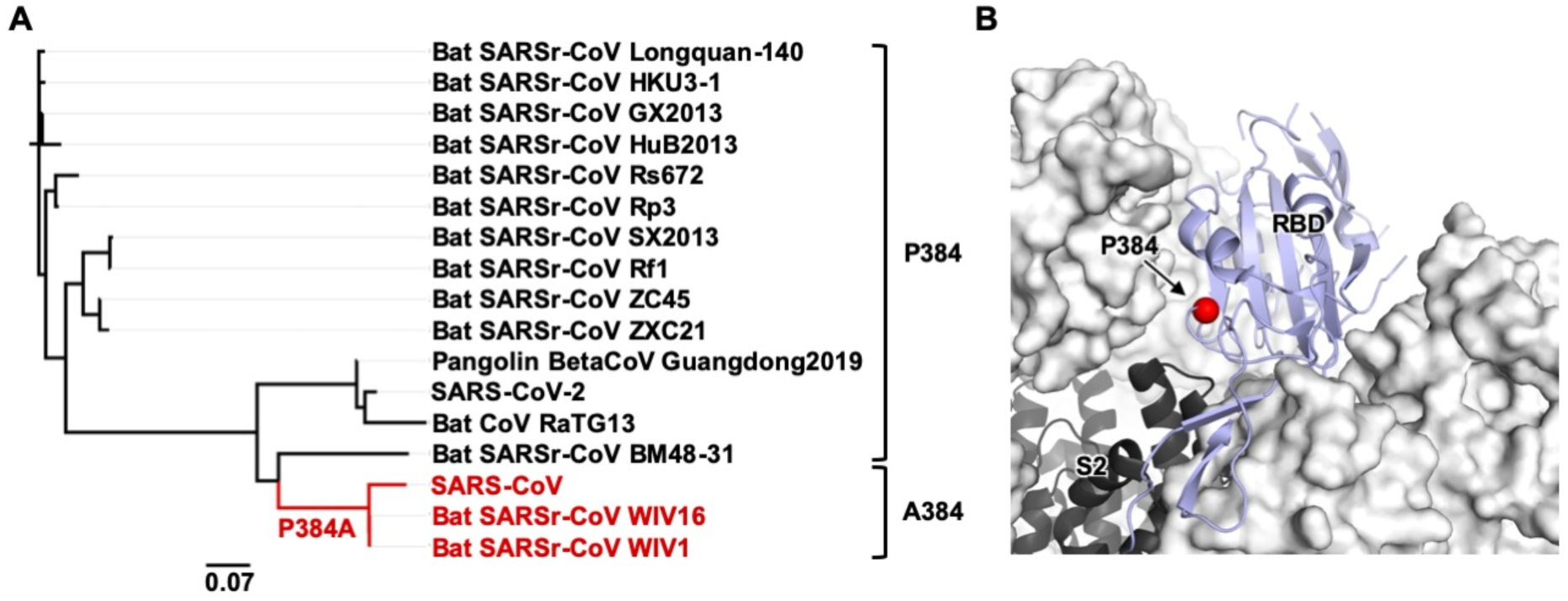
Sequence conservation and location of residue 384. **(A)** A phylogenetic tree was constructed based on the amino-acid sequences of RBDs from SARS-CoV-2, SARS-CoV, and SARS-related coronavirus (SARSr-CoV) strains. Branches corresponding to strains that have A384 are colored in red on the phylogenetic tree. Scale bar represents 0.07 amino-acid substitutions per position. **(B)** The location of P384 is shown on the SARS-CoV-2 S protein (PDB 6VXX [31]). S1 domain is represented by the white surface and the S2 domain by the black cartoon. The location of residue 384 is indicated by the red sphere on the RBD in the “down” conformation (blue cartoon).

### Crystal structure reveals the impact of P384A in CR3022 binding

We further determined the x-ray structure of SARS-CoV RBD in complex with CR3022 to 2.7 Å resolution (Figure 4A, Supplementary Table 1, and Supplementary Figure 1). The overall structure of CR3022 in complex with SARS-CoV RBD is similar to that with SARS-CoV-2 RBD [20] (Cα RMSD of 0.5 Å for 343 residues in the RBD and Fab variable domain, cf. fig. S2A and B of [20]) (Supplementary Figure 2). Nonetheless, the CR3022 elbow angles, which are distant from the antibody-antigen interface, differ in the two structures, as we mutated the elbow region (as described in [25]) of CR3022 to promote crystallization with SARS-CoV RBD. The conserved binding mode of CR3022 to SARS-CoV-2 RBD and SARS-CoV RBD indicates that the difference in binding affinity of CR3022 between SARS-CoV-2 RBD and SARS-CoV RBD is only due to a very subtle structural difference.

**Figure 4.**
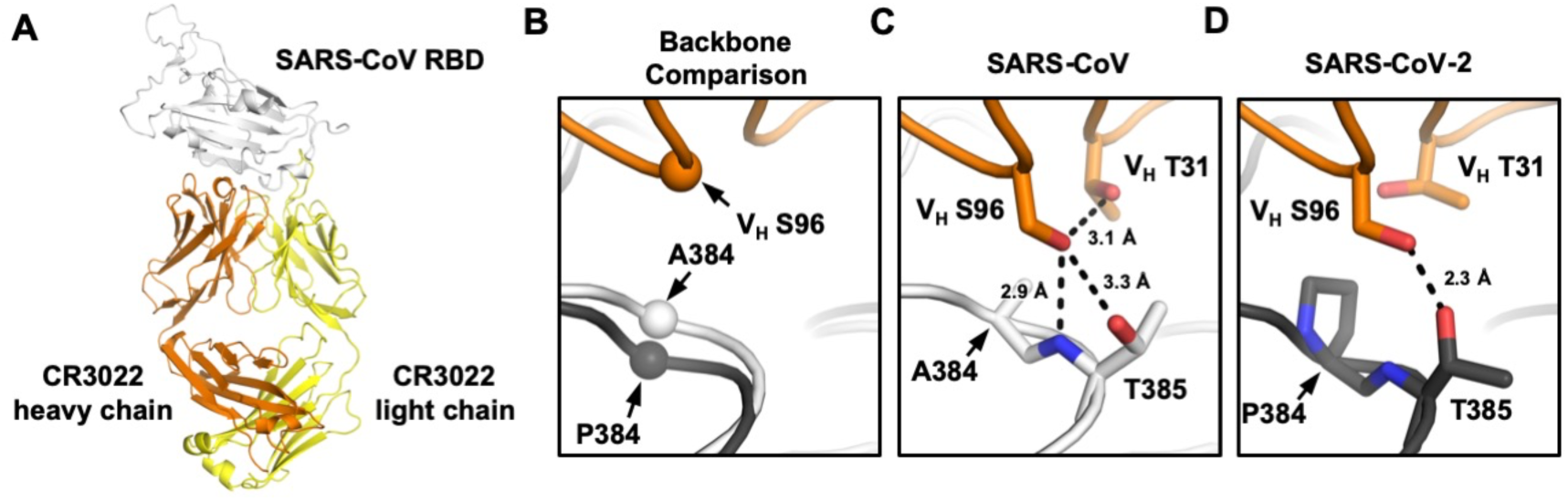
Crystal structure of CR3022 in complex with SARS-CoV RBD. **(A)** Crystal structure of CR3022 Fab in complex with SARS-CoV RBD. CR3022 heavy chain is colored in orange, CR3022 light chain in yellow, and SARS-CoV-2 RBD in light grey. **(B)** Structures of CR3022 in complex with SARS-CoV-2 RBD and with SARS-CoV RBD were aligned using the CR3022 heavy chain variable domain and the region around residue 384 is shown. Orange: CR3022 heavy chain. White: SARS-CoV RBD. Dark gray: SARS-CoV-2 RBD. The Cαs of S96 on CR3022 heavy chain, A384 on SARS-CoV RBD, and P384 on SARS-CoV-2 RBD are shown in sphere representation. **(C-D)** Interaction between CR3022 and residue 384 on **(C)** SARS-CoV RBD, and **(D)** SARS-CoV-2 RBD. Hydrogen bonds are represented by dashed lines.

To investigate how P384 and A384 lead to differential binding of CR3022, we compared the RBD structures from SARS-CoV and SAR-CoV-2 when bound with CR3022. The RBDs have a Cα RMSD of only 0.6 Å (0.7 Å for CR3022 epitope residues). At residue 384, the backbone of SARS-CoV-2 is further from CR3022, as compared to that of SARS-CoV (Figure 4B). This difference in backbone positioning (∼1.3 Å shift) affects the interaction of the RBD with CR3022 V_H_ S96, which is encoded by IGHD3-10 gene segment on CDR H3 [18, 20]. While CR3022 V_H_ S96 forms a hydrogen bond (H-bond) with the T385 side chain in both SARS-CoV-2 RBD and SARS-CoV RBD, it can form a second H-bond with the backbone amide of T385 in SARS-CoV RBD (Figure 4C), but not SARS-CoV-2 RBD (Figure 4D). In addition, CR3022 V_H_ S96 adopts different side-chain rotamers when binding to SARS-CoV-2 and to SARS-CoV. Consequently, V_H_ S96 can make an intramolecular H-bond with V_H_ T31 when CR3022 binds to SARS-CoV RBD (Figure 4C), but not to SARS-CoV-2 (Figure 4D). In summary, V_H_ S96 forms three H-bonds when CR3022 binds to SARS-CoV RBD, as compared to only one when CR3022 binds to SARS-CoV-2 RBD. This observation indicates why binding of CR3022 to the SARS-CoV RBD is energetically more favorable than to the SARS-CoV-2 RBD.

### CR3022-bound SARS-CoV S protein exhibits a rare three-up conformation

To understand the binding of CR3022 to the RBD in the context of the homotrimeric S protein, we previously proposed a structural model where CR3022 could only access its epitope on the S protein when at least two RBD are in the “up” conformation and the RBD is rotated relative to its unliganded structure [20]. To further evaluate and expand on this model, negative-stain electron microscopy (nsEM) was performed on CR3022 in complex with a stabilized version of the SARS-CoV homotrimeric S protein (Figure 5A, see Materials and Methods). The 3D nsEM reconstruction revealed that one SARS-CoV S protein could simultaneously bind to three CR3022 Fabs, with all three RBDs in the “up” conformation (Figure 5B). Consistent with the structural model that we previously proposed [20], the CR3022-bound RBD was indeed rotated compared to that in the unliganded S protein [26-28], such that, in this conformation, steric hinderance between CR3022 and the N-terminal domain (NTD) is minimized.

**Figure 5.**
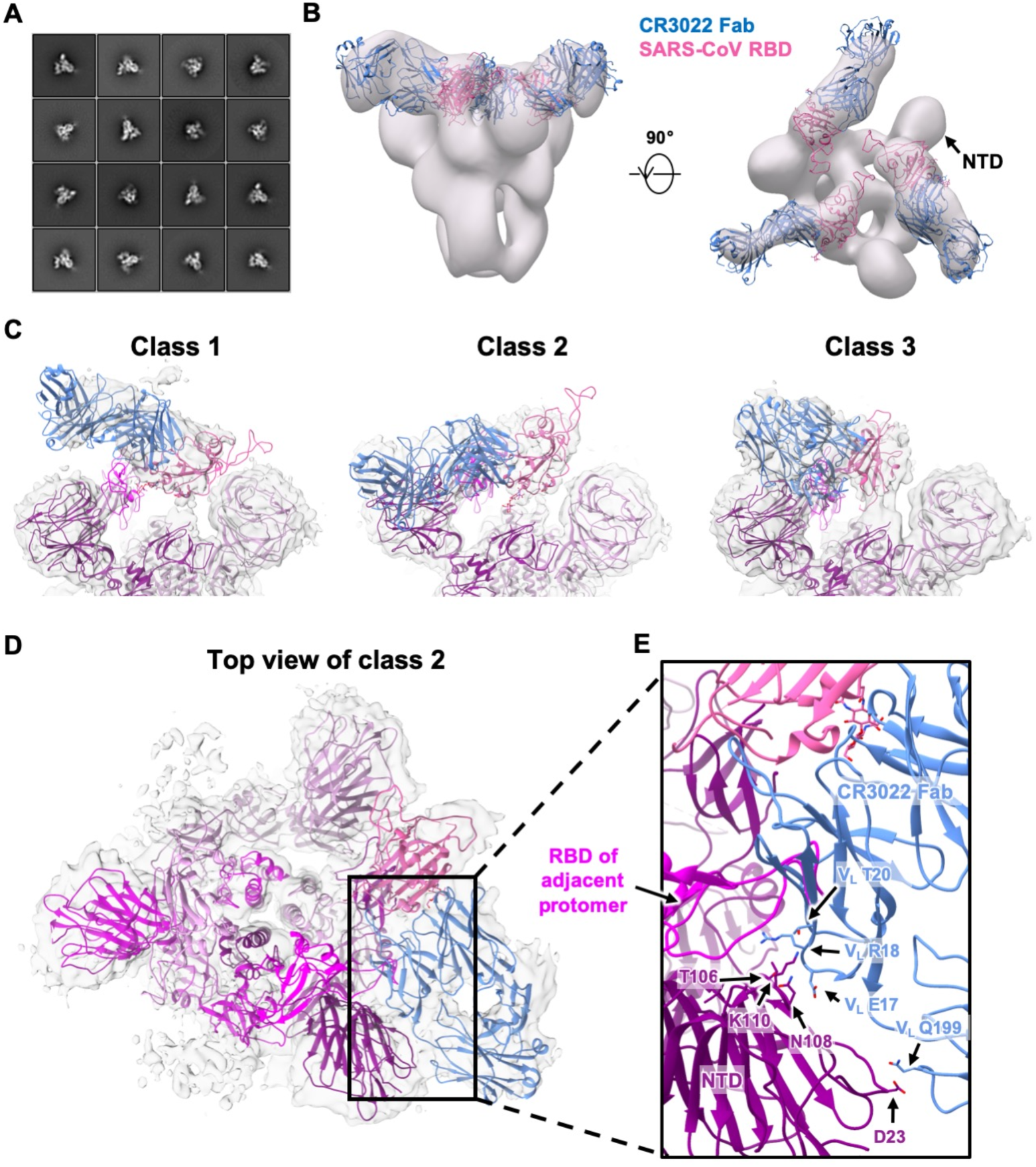
Negative-stain EM and cryo-EM analysis of SARS spike bound to CR3022 Fab. **(A)** Representative 2D nsEM class averages of the trimeric SARS-CoV spike glycoprotein complexed with three CR3022 Fabs. **(B)** Side and top view of the 3D reconstruction showing CR3022 Fabs bound to all 3 RBDs on the SARS-CoV spike. The SARS-CoV RBD-CR3022 complex from the crystal structure is fitted into the nsEM density with the RBD shown in pink and CR3022 Fab in blue. **(C)** Side views of the B-factor-sharpened cryo-EM maps (transparent gray surface representation) representing three different classes of SARS spike with CR3022 Fab with different RBD-Fab orientations. While four different classes were identified, only three classes are shown here because classes 2 and 4 are very similar (Supplementary Figure 4). The RBD-Fab complex model is fit into the densities with the RBDs shown in pink and CR3022 Fabs represented in blue. The atomic model of the apo SARS-CoV spike (PDB 6ACD) [35] is also fit into density with one RBD removed for clarity. The protomers are colored in purple, magenta and deep magenta. **(D)** Top view of the class 2 cryo-EM map depicting potential quaternary contacts between the RBD-bound Fab and the spike NTD in this conformation. In this RBD-Fab conformation, the Fab would clash with the “down” RBD of the adjacent protomer (magenta) and, therefore, the adjacent RBD can only exist in an “up” conformation. **(E)** A close-up view of the Fab-spike interface showing the superimposition of CR3022 Fab and adjacent RBD. The residues that can contribute to quaternary interactions between CR3022 light chain and the NTD in two of the four classes (2 and 4) are shown.

While our results here demonstrate that CR3022 Fab could form a stable complex with SARS-CoV S protein in a prefusion conformation, a recent study reported that prefusion SARS-CoV-2 S protein fell apart upon binding to CR3022 Fab as indicated by cryo-EM [29]. It should be noted that the three-up conformation is much more rarely observed than the other RBD conformations (all-down, one-up, and two-up) in SARS-CoV by cryo-EM [26-28], or SARS-CoV-2 by cryo-EM [30-32] and cryo-electron tomography [33, 34], and could relate to differences in the stability of S trimers in SARS-CoV versus SARS CoV-2 when CR3022 is bound. Further studies will be required to investigate whether such a difference between SARS-CoV-2 and SARS-CoV is related to stability differences in the recombinant spike proteins, or to different dynamics of the RBD on the virus or infected cells.

### RBD flexibility and quaternary interactions in CR3022-bound SARS-CoV S protein

To address some of these issues, we performed cryo-EM analysis to interrogate the binding of CR3022 to SARS-CoV S protein at higher resolution (Supplementary Figure 3 and Supplementary Table 2). Focused 3D classification yielded 4 different structural classes with classes 2 and 4 being nearly identical at the given resolution (Figure 5C and Supplementary Figure 4). Class 3 is the most similar to the model from nsEM, although the total particle number for classes 2 and 4 together exceed that for class 3 (Supplementary Figure 4). In contrast, class 1 is the least represented. In classes 2 and 4, CR3022 also appears to make quaternary contacts with the NTD, as suggested by well-defined density in the CR3022-NTD interface (Figure 5C). The moderate resolution (6 to 7 Å) of the reconstructions precludes atomic-level descriptions, but the framework region of the CR3022 light chain in classes 2 and 4 is in close proximity to a loop region in NTD corresponding to residues 106-110. In addition, the constant region of CR3022 appears to contact residue D23 of NTD. Another important observation is that the Fab in class 2 and 4 would clash with the adjacent RBD if it were in the “down” conformation. So, for the Fab to exist in this quaternary conformation, the adjacent RBD has to be in the “up” conformation. To evaluate the different dispositions of the RBD in these structures, we compared the cryo-EM structure of an apo form of the SARS-CoV S protein where one RBD is the “up” conformation (PDB 6ACD) [35]. The RBD in classes 1 to 4 are rotated by 84.1°, 54.3°, -54.7°, and 53.7°, respectively, relative to the apo one-up conformation (see Materials and Methods). Furthermore, the CR3022-bound RBD in class 2 and 4 is more open than in the apo form (Supplementary Figure 5), demonstrating the rotational flexibility of the RBD. In fact, RBD conformational flexibility has also been noted in an ACE2-bound SARS-CoV S protein. Three different dispositions (1 to 3) of the RBD were observed in ACE2-bound SARS-CoV S protein with RBD tilts relative to horizontal top surface of the S trimer of 51.2°, 73.3° and 111.6° compared to 68.9° for the apo one-up structure [35]. Our classes 2 and 4 appear to be somewhat intermediate between dispositions 2 and 3 (Supplementary Figure 6), whereas the other classes differ from the RBD dispositions in the ACE2-bound SARS-CoV S structures. However, despite the flexibility of CR3022-bound RBD, bivalent binding of CR3022 to S protein does not seem to occur on the virus surface since an IgG avidity effect was not observed in the neutralization assay (see above, Figure 2). Overall, these structural analyses indicate that RBD rotational flexibility and acquisition of quaternary interactions can play an important role in CR3022 interaction with the S protein. CR3022 adds to the growing list of neutralization antibodies that can utilize quaternary interactions for binding to the S protein [12, 36].

## DISCUSSION

While it is now known that SARS-CoV and SARS-CoV-2 differ in antigenicity despite relatively high sequence conservation [1, 3, 4, 14], there is a paucity of understanding of the underlying molecular determinants of these antigenic changes and the structural consequences of these differences. Through structural analysis of the CR3022-RBD complex and mutagenesis experiments, we show that a single amino-acid substitution at residue 384 contributes to an important antigenic difference in a highly conserved (neutralizing) epitope between SARS-CoV-2 and SARS-CoV.

While CR3022 cannot neutralize SARS-CoV-2 WT in almost all studies [3, 5, 20, 22], it can neutralize the SARS-CoV-2 P384A mutant. The K_D_ of CR3022 Fab to SARS-CoV-2 WT RBD is 68 nM, whereas to SARS-CoV-2 P384A RBD is 1 nM (Figure 1B-C), indicating that the affinity threshold for neutralization of SARS-CoV-2 to this epitope is in the low nM range. However, despite having a low nM affinity to SARS-CoV-2 P384A RBD, CR3022 only weakly neutralizes SARS-CoV-2 P384A with an IC_50_ of 3.2 μg/ml and SARS-CoV with an IC_50_ of 5.2 μg/ml. In contrast, antibodies with similar or less Fab binding affinity to other RBD epitopes, such as the receptor binding site, can neutralize SARS-CoV-2 much more efficiently. For example, previously characterized SARS-CoV-2 antibodies CC12.1 and CC12.3, which have a K_D_ to SARS-CoV-2 RBD of 17 nM and 14 nM respectively, neutralize SARS-CoV-2 at an IC_50_ of ∼20 ng/ml [3, 37]. Of note, the K_D_ and IC_50_ of CC12.1 and CC12.3 were measured in the same manner as this study. The lack of correlation between affinity and neutralizing activity is therefore not due to the difference in the assays between studies. In fact, a previous study also demonstrated a lack of correlation between RBD binding and neutralization for monoclonal antibodies [3]. Together, these observations suggest that the affinity threshold for SARS-CoV-2 neutralization by RBD-targeting antibodies may be epitope dependent. The difference in affinity threshold for different epitopes is also likely to be related not only in the ability to block ACE2-binding [3, 38], but also in antibody avidity where bivalent binding can cross-link different RBD domains on the same or different spikes and, hence, substantially enhance binding and neutralization [23].

Given the scale of the outbreak, SARS-CoV-2 may persist and circulate in humans for years to come [39]. A number of SARS-CoV-2 vaccine candidates are currently under clinical trials (https://clinicaltrials.gov/ct2/who_table) [40], which offer a potential solution to alleviate the global health and socio-economic devastation bought by SARS-CoV-2. However, whether SARS-CoV-2 can escape vaccine-induced immunity through antigenic drift remains to be determined, although escape mutations to many monoclonal antibodies have been tested *in vitro* [2]. Identification of the key residues that are responsible for differences in antigenicity among SARS-CoV-2, SARS-CoV, and possibly other SARS-related viruses, should provide a starting point to understand the potential for antigenic drift in SARS-like coronaviruses. The ongoing efforts in SARS-CoV-2 antibody discovery and structural characterization will therefore advance our molecular understanding of antigenic variation in SARS-like CoVs, and consequences for vaccine and therapeutic design, especially to cross-neutralizing epitopes.

## ACKNOWLEDGEMENTS

We thank Henry Tien for technical support with the crystallization robot, Jeanne Matteson for contribution to mammalian cell culture, Wenli Yu to insect cell culture, Robyn Stanfield for assistance in data collection, and Chris Mok for pilot testing of the pseudovirus assay. We are grateful to the staff of Stanford Synchrotron Radiation Laboratory (SSRL) Beamline 12-2 for assistance. This work was supported by NIH R00 AI139445 (N.C.W.), NIH R01 AI073148 (D.N.), the Bill and Melinda Gates Foundation OPP1170236 (A.B.W. and I.A.W.) and NIH CHAVD UM1 AI44462 (A.B.W. and I.A.W.). Use of the SSRL, SLAC National Accelerator Laboratory, is supported by the U.S. Department of Energy, Office of Science, Office of Basic Energy Sciences under Contract No. DE-AC02–76SF00515. The SSRL Structural Molecular Biology Program is supported by the DOE Office of Biological and Environmental Research, and by the National Institutes of Health, National Institute of General Medical Sciences (including P41GM103393).

## AUTHOR CONTRIBUTIONS

N.C.W., M.Y., D.H., S.B., A.B.W and I.A.W. conceived and designed the study. N.C.W., M.Y., C.C.D.L. and S.B. expressed and purified the proteins. M.Y. performed biolayer interferometry binding assays. N.C.W. and M.Y. performed the crystallization experiment and X.Z. collected the X-ray data. M.Y. determined and refined the X-ray structures. S.B. and H.L.T. performed the negative-stain electron microscopy. D.H., L.P., L.Y. and D.N. performed the pseudovirus neutralization assay. N.C.W., M.Y., D.H., S.B., A.B.W. and I.A.W. analyzed the data. N.C.W., M.Y. and I.A.W. wrote the paper and all authors reviewed and edited the paper.

## MATERIALS AND METHODS

### Expression and purification of SARS-CoV RBD

RBD (residues: 306-527) of the SARS-CoV spike (S) protein (GenBank: ABF65836.1) was fused with an N-terminal gp67 signal peptide and a C-terminal His_6_ tag, and cloned into a customized pFastBac vector [41]. Recombinant bacmid DNA was generated using the Bac-to-Bac system (Thermo Fisher Scientific). Baculovirus was generated by transfecting purified bacmid DNA into Sf9 cells using FuGENE HD (Promega), and subsequently used to infect suspension cultures of High Five cells (Thermo Fisher Scientific) at an MOI of 5 to 10. Infected High Five cells were incubated at 28 °C with shaking at 110 r.p.m. for 72 h for protein expression. The supernatant was then concentrated using a 10 kDa MW cutoff Centramate cassette (Pall Corporation). SARS-CoV RBD protein was purified by Ni-NTA, followed by size exclusion chromatography, and buffer exchanged into 20 mM Tris-HCl pH 7.4 and 150 mM NaCl.

### Expression and purification of SARS-CoV spike

The SARS-CoV spike construct (Tor2 strain) for recombinant spike protein expression contains the mammalian-codon-optimized gene encoding residues 1-1190 of the spike followed by a C-terminal T4 fibritin trimerization domain, a HRV3C cleavage site, 8x-His tag and a Twin-strep tags subcloned into the eukaryotic-expression vector pαH. Residues at 968 and 969 were replaced by prolines for generating stable spike proteins as described previously [28]. The spike plasmid was transfected into FreeStyle 293F cells and cultures were harvested at 6-day post-transfection. Proteins were purified from the supernatants on His-Complete columns using a 250 mM imidazole elution buffer. The elution was buffer exchanged to Tris-NaCl buffer (25 mM Tris, 500 mM NaCl, pH 7.4) before further purification using Superose 6 increase 10/300 column (GE Healthcare). Protein fractions corresponding to the trimeric spike proteins were collected and concentrated.

### Expression and purification of CR3022 Fab

The CR3022 Fab heavy (GenBank: DQ168569.1) and light (GenBank: DQ168570.1) chains were cloned into phCMV3. The plasmids were transiently co-transfected into Expi293F cells at a ratio of 2:1 (HC:LC) using ExpiFectamine™ 293 Reagent (Thermo Fisher Scientific) according to the manufacturer” s instructions. The supernatant was collected at 7 days post-transfection. The Fab was purified with a CaptureSelect™ CH1-XL Pre-packed Column (Thermo Fisher Scientific) followed by size exclusion chromatography. For crystallization, a VSRRLP variant of the elbow region was used to reduce the conformational flexibility between the constant and variable domains [25].

### Crystallization and structural determination

Purified CR3022 Fab with a VSRRLP modification in the elbow region and SARS-CoV RBD were mixed at a molar ratio of 1:1 and incubated overnight at 4°C. The complex (7.5 mg/ml) was screened for crystallization using the 384 conditions of the JCSG Core Suite (Qiagen) on our custom-designed robotic CrystalMation system (Rigaku) at Scripps Research by the vapor diffusion method in sitting drops containing 0.1 μl of protein and 0.1 μl of reservoir solution. Optimized crystals were then grown in 2 M sodium chloride and 10% PEG 6000 at 4°C. Crystals were grown for 7 days and then flash cooled in liquid nitrogen. Diffraction data were collected at cryogenic temperature (100 K) at Stanford Synchrotron Radiation Lightsource (SSRL) beamline 12-2 with a wavelength of 1.033 Å, and processed with HKL2000 [42]. Structures were solved by molecular replacement using PHASER [43] with PDB 6W41 for CR3022 Fab [20] and PDB 2AJF for SARS-CoV RBD [44]. Iterative model building and refinement were carried out in COOT [45] and PHENIX [46], respectively. Ramachandran statistics were calculated using MolProbity [47].

### Negative-stain electron microscopy

Six molar excess of CR3022 Fab (unmodified) was added to SARS-CoV spike protein 1 hour prior to direct deposition onto carbon-coated 400-mesh copper grids. The grids were stained with 2 % (w/v) uranyl-formate for 90 seconds immediately following sample application. Grids were imaged on Tecnai T12 Spirit at 120 keV with a 4k x 4k Eagle CCD. Micrographs were collected using Leginon [48] and images were transferred to Appion [49] for particle picking using a difference-of-Gaussians picker (DoG-picker) [50] and generation of particle stacks. Particle stacks were further transferred to Relion [51] for 2D classification followed by 3D classification to select good classes. Select 3D classes were auto-refined on Relion and used for making figures using UCSF Chimera [52].

### Cryo-EM sample preparation

SARS-CoV spike protein was incubated with six molar excess of CR3022 Fab for 2 h. 3.5 µL of the complex (0.9 mg/ml) was mixed with 0.5 µL of 0.04 mM lauryl maltose neopentyl glycol (LMNG) solution immediately before sample deposition onto a 1.2/1.3 300-Gold grid (EMS). The grids were plasma cleaned for 7 seconds using a Gatan Solarus 950 Plasma system prior to sample deposition. Following sample application, grids were blotted for 3 seconds before being vitrified in liquid ethane using a Vitrobot Mark IV (Thermo Fisher).

### Cryo-EM data collection and processing

Data collection was performed using a Talos Arctica TEM at 200 kV with a Gatan K2 Summit detector at a magnification of 36,000x, resulting in a 1.15 Å pixel size. Total exposure was split into 250 ms frames with a total cumulative dose of ∼50 e^-^/Å^2^. Micrographs were collected through Leginon software at a nominal defocus range of -0.4 µm to -1.6 µm and MotionCor2 was used for alignment and dose weighting of the frames [48, 53]. Micrographs were transferred to CryoSPARC 2.9 for further processing [54]. CTF estimations were performed using GCTF and micrographs were selected using the Curate Exposures tool in CryoSPARC based on their CTF resolution estimates (cutoff 5 Å) for downstream particle picking, extraction and iterative rounds of 2D classification and selection [55]. Particles selected from 2D classes were transferred to Relion 3.1 for direct C3 refinement, symmetry expansion of particles and iterative rounds of 3D focused classification using spherical masks around the RBD and Fab [51]. Final subsets of clean particles from 4 different classes were each refined with C1 symmetry. Figures were generated using UCSF Chimera and UCSF Chimera X [52].

### Calculation of rotation angles

Comparisons of subunit rotation angles among different structures were performed with a software “Superpose” in the CCP4 package [56, 57]. For each classified conformation, the Cα atoms of the RBD domain are superimposed to the equivalent atoms of the RBD in “up” -conformation in a previously reported spike trimer cryoEM structure (PDB 6ACD) [35]. The rotation matrices generated by superposing each pair of structures with “Superpose” were adopted to calculate the subunit rotation angle following the equation shown as below:

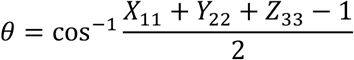

where *θ* is the subunit rotation angle, *X*_11_, *Y*_22_, and *Z*_33_ represent the *X*_11_, *Y*_22_, and *Z*_33_ values in the rotation matrix calculated for the superpose.

### Biolayer interferometry binding assay

Binding assays were performed by biolayer interferometry (BLI) using an Octet Red instrument (FortéBio) as described previously [58]. Briefly, His_6_-tagged SARS-CoV RBD proteins at 20 to 100 μg/ml in 1x kinetics buffer (1x PBS, pH 7.4, 0.01% BSA and 0.002% Tween 20) were loaded onto Anti-Penta-HIS (HIS1K) biosensors and incubated with the indicated concentrations of CR3022 Fab. The assay consisted of five steps: 1) baseline: 60 s with 1x kinetics buffer; 2) loading: 300 s with His_6_-tagged S or RBD proteins; 3) baseline: 60 s with 1x kinetics buffer; 4) association: 120 s with samples (Fab or IgG); and 5) dissociation: 120 s with 1x kinetics buffer. For estimating the exact K_D_, a 1:1 binding model was used.

### Pseudovirus neutralization assay

Pseudovirus preparation and assay were performed as previously described [3]. Briefly, MLV-gag/pol and MLV-CMV plasmids was co-transfected into HEK293T cells along with full-length or P384A SARS-CoV-2 spike plasmids using Lipofectamine 2000 to produce pseudoviruses competent for single-round infection. The supernatant containing MLV-pseudotyped viral particles was collected at 48 hours post transfection, aliquoted and frozen at -80°C until used. For each well in a 96-well half-area plate, 25 μl of virus was immediately mixed with 25 μl of serially diluted IgG or Fab, and incubated for 1 hour at 37°C. For each well, 10,000 HeLa-hACE2 cells in 50 μl of media supplemented with 20 μg/ml dextran were added to the antibody-virus mixture. The 96-well half-area plate was then incubated at 37°C. At 42 to 48 hours post-infection, HeLa-hACE2 cells were lysed using 1x luciferase lysis buffer (25 mM Gly-Gly pH 7.8, 15 mM MgSO_4_, 4 mM EGTA, and 1% Triton X-100). Luciferase intensity was then measured using Bright-Glo Luciferase Assay System (Promega) according to the manufacturer” s instructions. Percentage of neutralization was calculated using the following equation:

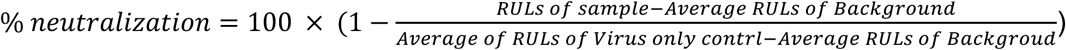

## DATA AVAILABILITY

The X-ray coordinates and structure factors have been deposited to the RCSB Protein Data Bank under accession code: 7JN5. The EM maps will be deposited in the Electron Microscopy Data Bank (EMDB).

---

**Supplementary Table 1.** X-ray data collection and refinement statistics.

|  |  |
| --- | --- |
| **Data collection** |  |
| Beamline | SSRL 12-2 |
| Wavelength (Å) | 0.97946 |
| Space group | C 1 2 1 |
| Unit cell parameters (Å and °) | a=265.7, b=59.9, c=51.7, β=99.8 |
| Resolution (Å) | 50.0–2.70 (2.76–2.70)<sup>a</sup> |
| Unique reflections | 21,547 (2,021)<sup>a</sup> |
| Redundancy | 6.7 (5.5)<sup>a</sup> |
| Completeness (%) | 100.0 (100.0)<sup>a</sup> |
| <I/σ<sub>I</sub>> | 14.7 (1.0)<sup>a</sup> |
| R<sub>sym</sub><sup>b</sup> (%) | 9.2 (86.1)<sup>a</sup> |
| R<sub>pim</sub><sup>b</sup> (%) | 5.4 (54.8)<sup>a</sup> |
| CC<sub>1/2</sub><sup>c</sup> (%) | 99.4 (74.4)<sup>a</sup> |
| **Refinement statistics** |  |
| Resolution (Å) | 45.0–2.70 |
| Reflections (work) | 21,501 |
| Reflections (test) | 1,011 |
| R<sub>cryst</sub><sup>d</sup> / R<sub>free</sub><sup>e</sup> (%) | 22.2 / 27.6 |
| No. of atoms | 4,872 |
| Macromolecules | 4,795 |
| Glycans | 42 |
| Solvent | 30 |
| Average B-value (Å<sup>2</sup>) | 80 |
| Macromolecules | 80 |
| RBD | 104 |
| Fab | 70 |
| Glycans | 30 |
| Solvent | 60 |
| Wilson B-value (Å<sup>2</sup>) | 64 |
| **RMSD from ideal geometry** |  |
| Bond length (Å) | 0.005 |
| Bond angle (°) | 1.17 |
| **Ramachandran statistics (%)** |  |
| Favored | 95.6 |
| Outliers | 0.16 |
| **PDB code** | **7JN5** |

<sup>a</sup> Numbers in parentheses refer to the highest resolution shell.
<sup>b</sup> $R_{\text{sym}} = \sum_{hkl} \sum_i |I_{hkl,i} - \langle I_{hkl} \rangle| / \sum_{hkl} \sum_i I_{hkl,i}$ and $R_{\text{pim}} = \sum_{hkl} (1/(n-1))^{1/2} \sum_i |I_{hkl,i} - \langle I_{hkl} \rangle| / \sum_{hkl} \sum_i I_{hkl,i}$ , where $I_{hkl,i}$ is the scaled intensity of the $i^{\text{th}}$ measurement of reflection $h, k, l$ , $\langle I_{hkl} \rangle$ is the average intensity for that reflection, and $n$ is the redundancy.
<sup>c</sup> CC<sub>1/2</sub> = Pearson correlation coefficient between two random half datasets.
<sup>d</sup> $R_{\text{cryst}} = \sum_{hkl} |F_o - F_c| / \sum_{hkl} |F_o| \times 100$ , where $F_o$ and $F_c$ are the observed and calculated structure factors, respectively.
<sup>e</sup> $R_{\text{free}}$ was calculated as for $R_{\text{cryst}}$ , but on a test set comprising 5% of the data excluded from refinement.

**Supplementary Table 2.**
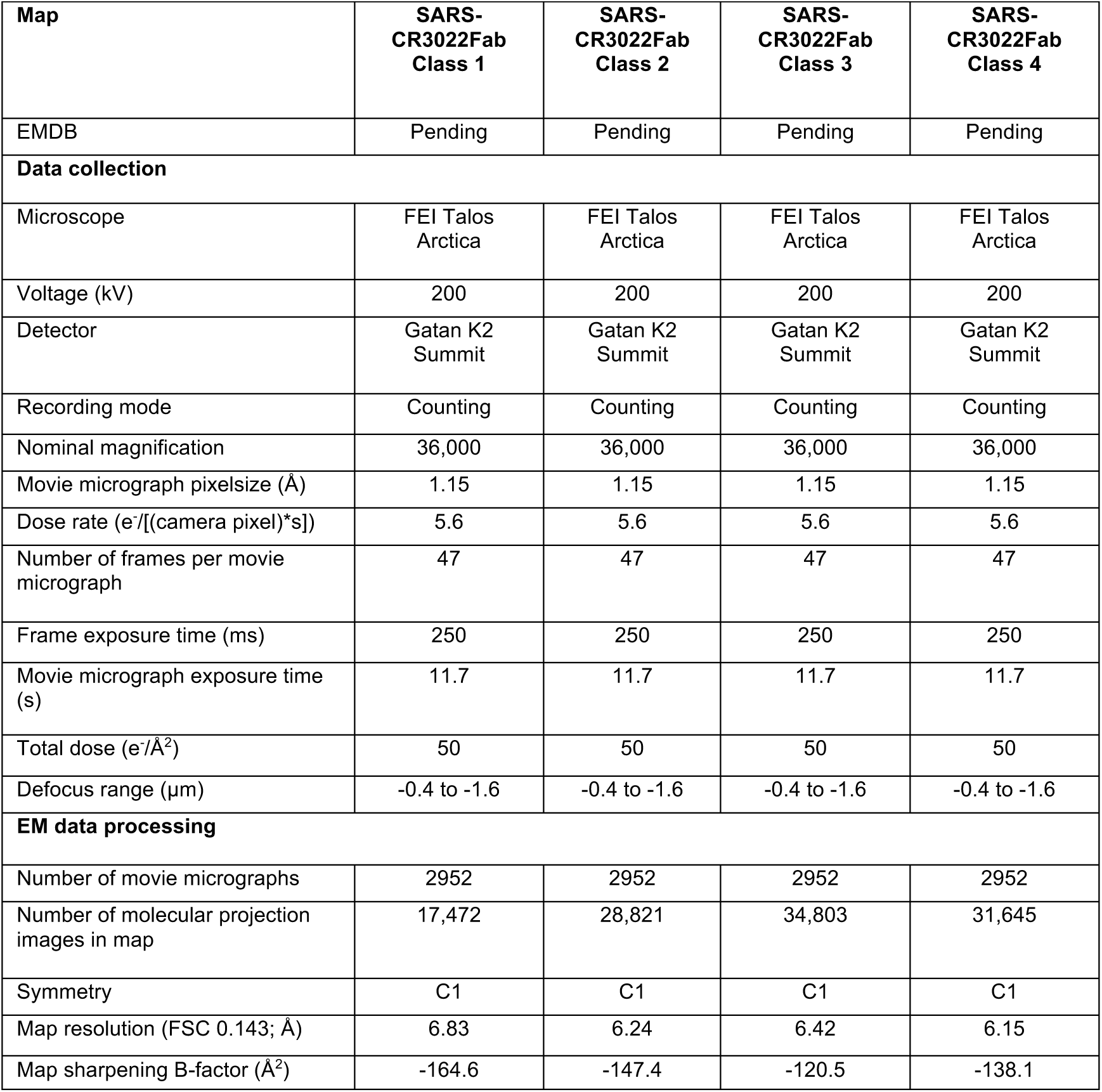
Cryo-EM data collection and refinement statistics.

**Supplementary Figure 1.**
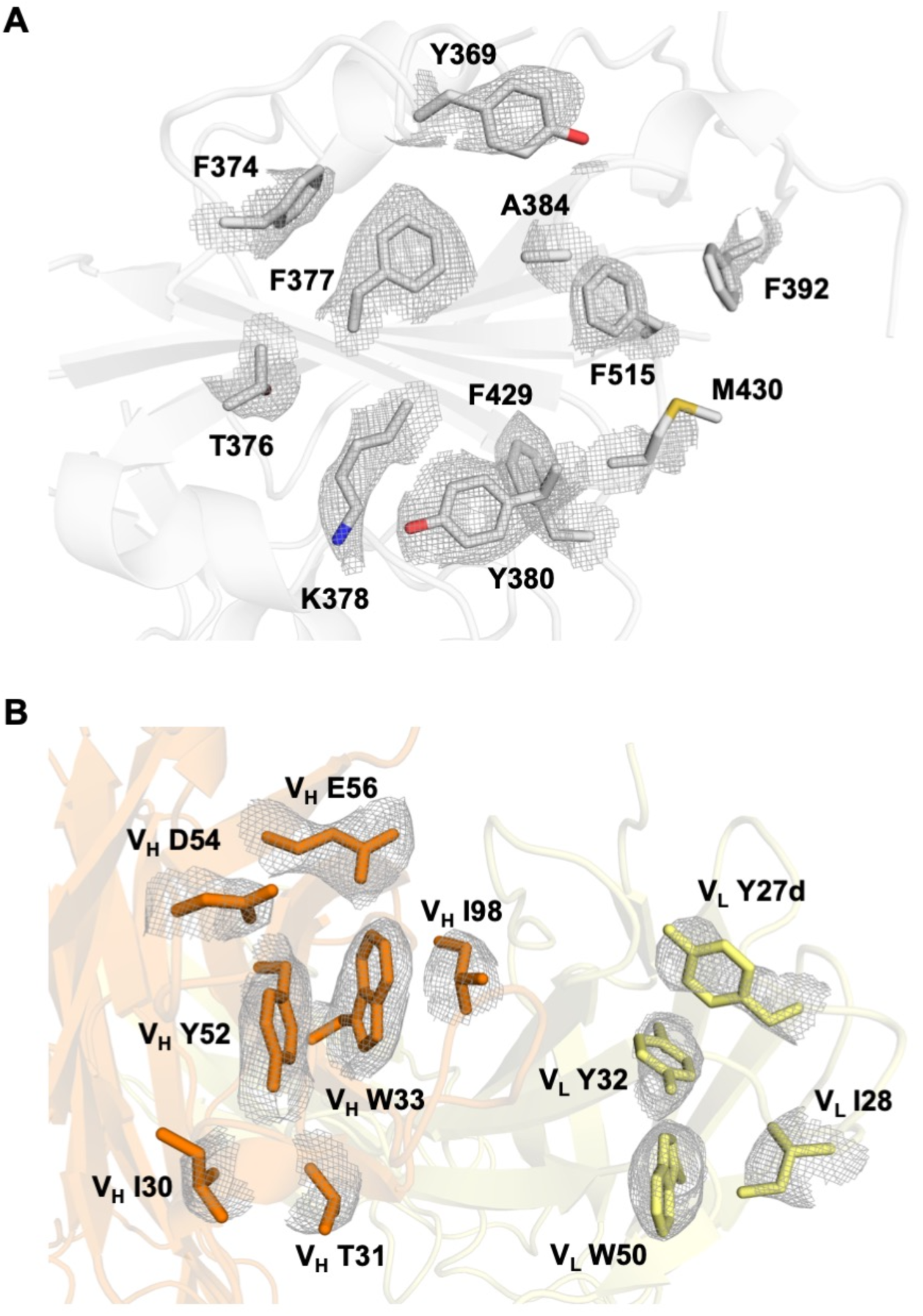
X-ray electron density maps for epitope and paratope regions of SARS CoV RBD with Fab CR3022. **(A)** Final 2Fo-Fc electron density maps for the side chains in the epitope region of SARS-CoV-2 contoured at 1 σ. **(B)** Final 2Fo-Fc electron density maps for the paratope region of CR3022 contoured at 1 σ. The heavy chain is colored in orange, and light chain in yellow. Epitope and paratope residues are labeled.

**Supplementary Figure 2.**
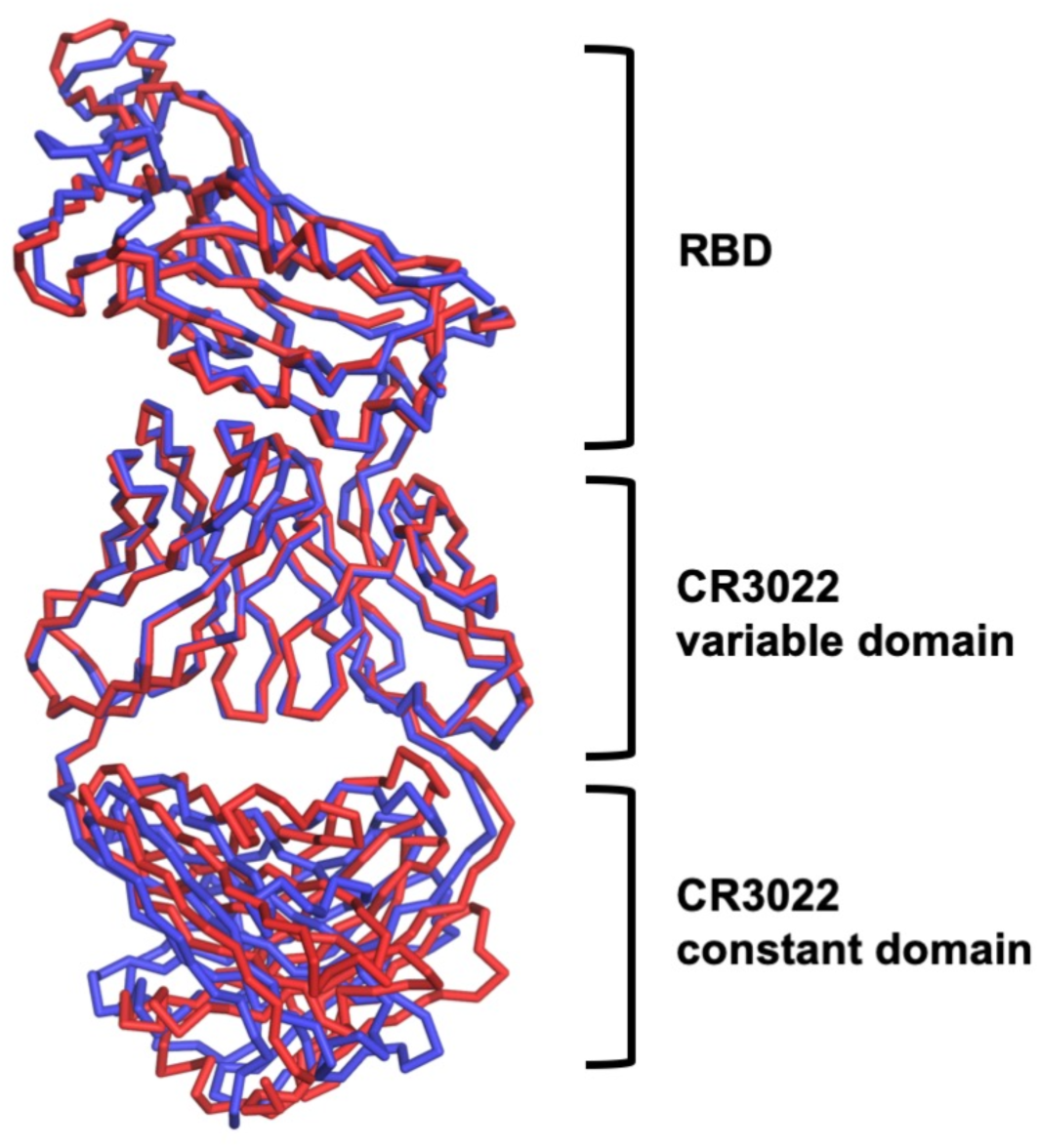
Structural alignment of CR3022-bound SARS-CoV RBD and CR3022-bound SARS-CoV-2 RBD. Structure of CR3022 in complex with SARS-CoV RBD (this study) is aligned to that with SARS-CoV-2 RBD (PDB 6W41). Structural alignment was performed using CR3022 heavy chain variable domain. Red: CR3022 in complex with SARS-CoV RBD. Blue: CR3022 in complex with SARS-CoV-2 RBD.

**Supplementary Figure 3.**
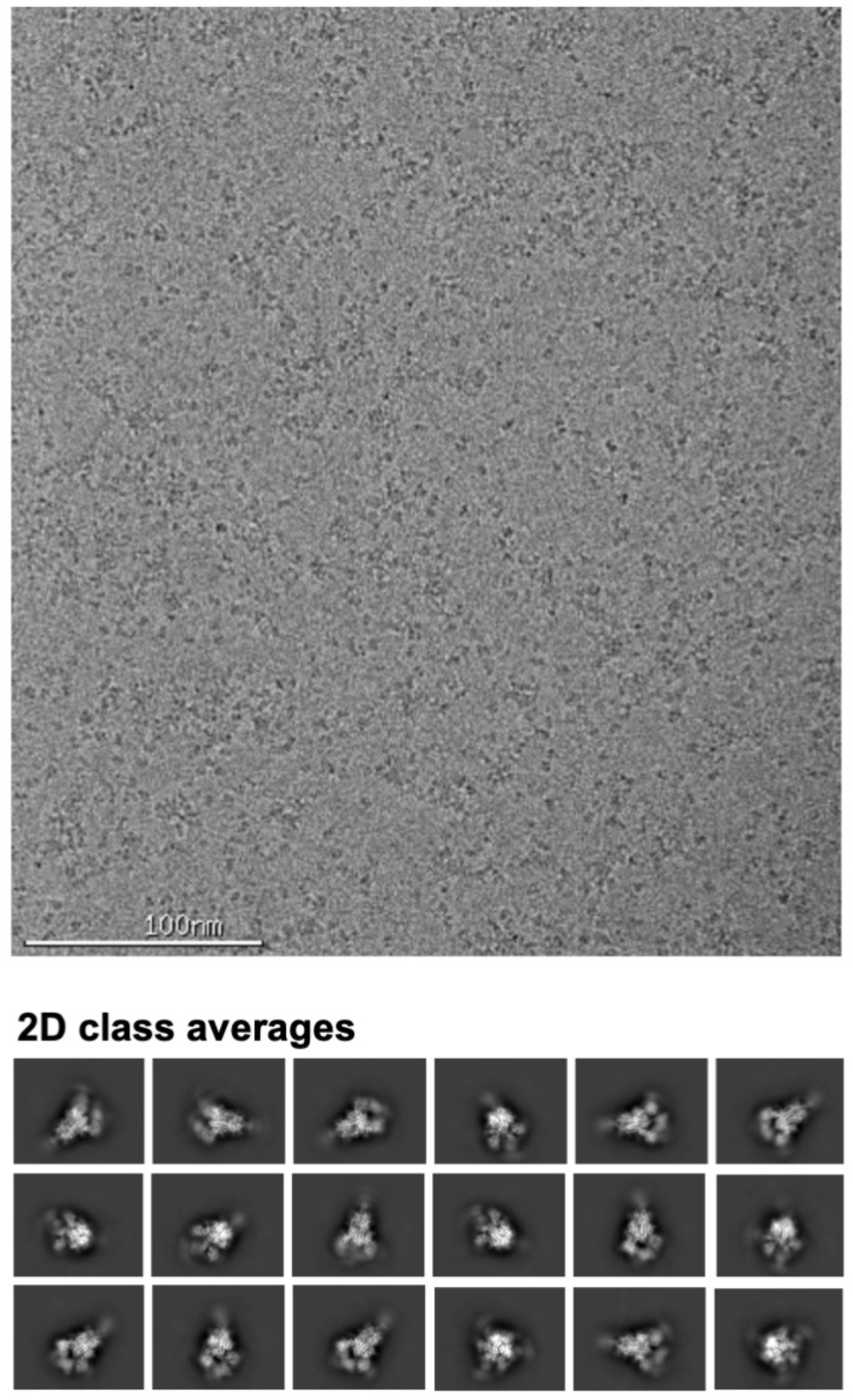
Representative cryo-electron micrograph and 2D class averages of the SARS-CoV spike in complex with CR3022 Fab. The top panel shows a representative cryo-electron micrograph of the SARS-CoV spike complexed with CR3022 Fab, whereas the bottom panels show the 2D class averages.

**Supplementary Figure 4.**
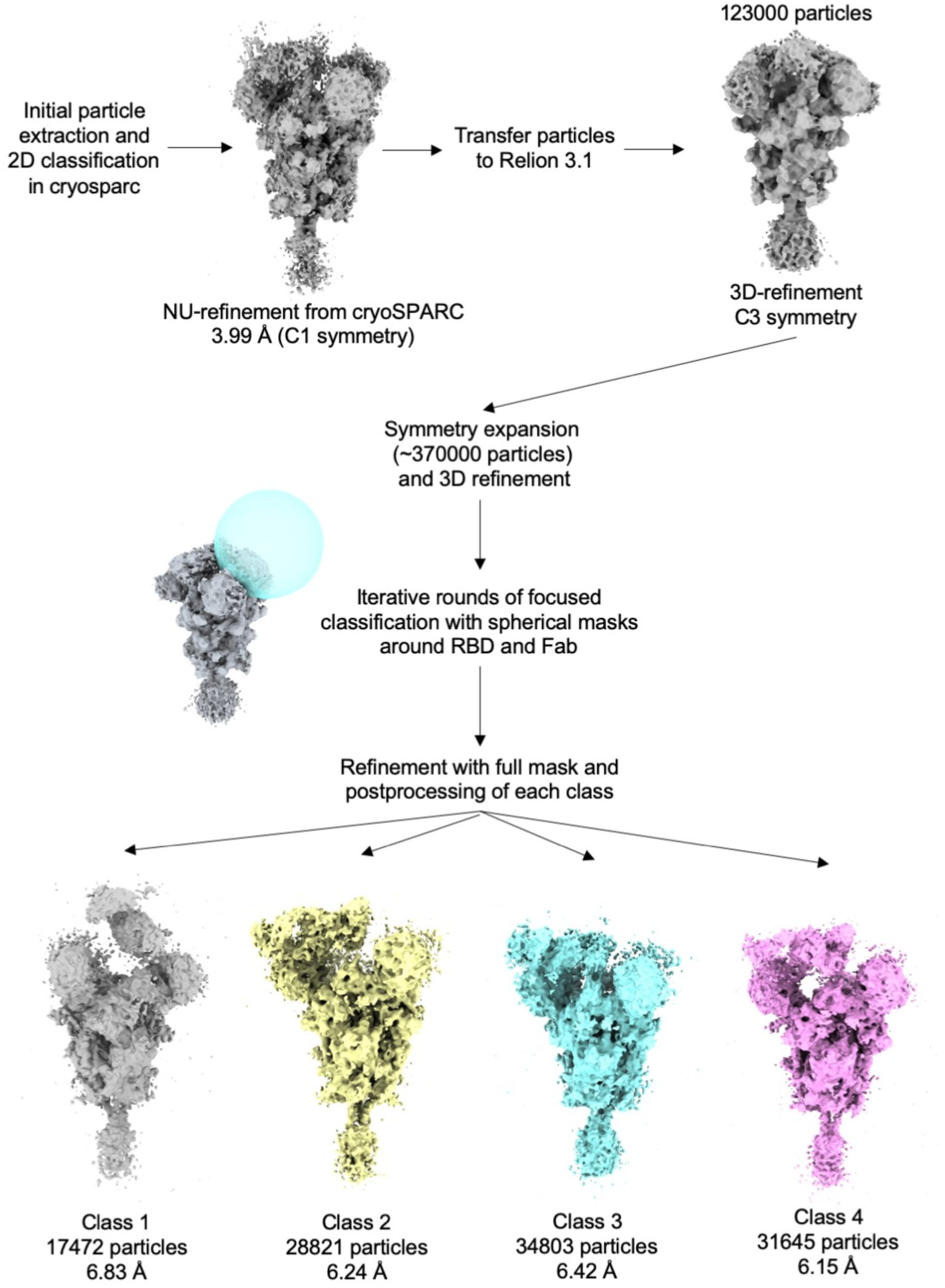
Workflow for cryo-EM data processing. Four 3D class averages of complex of the SARS-CoV spike and CR3022 were found during data processing.

**Supplementary Figure 5.**
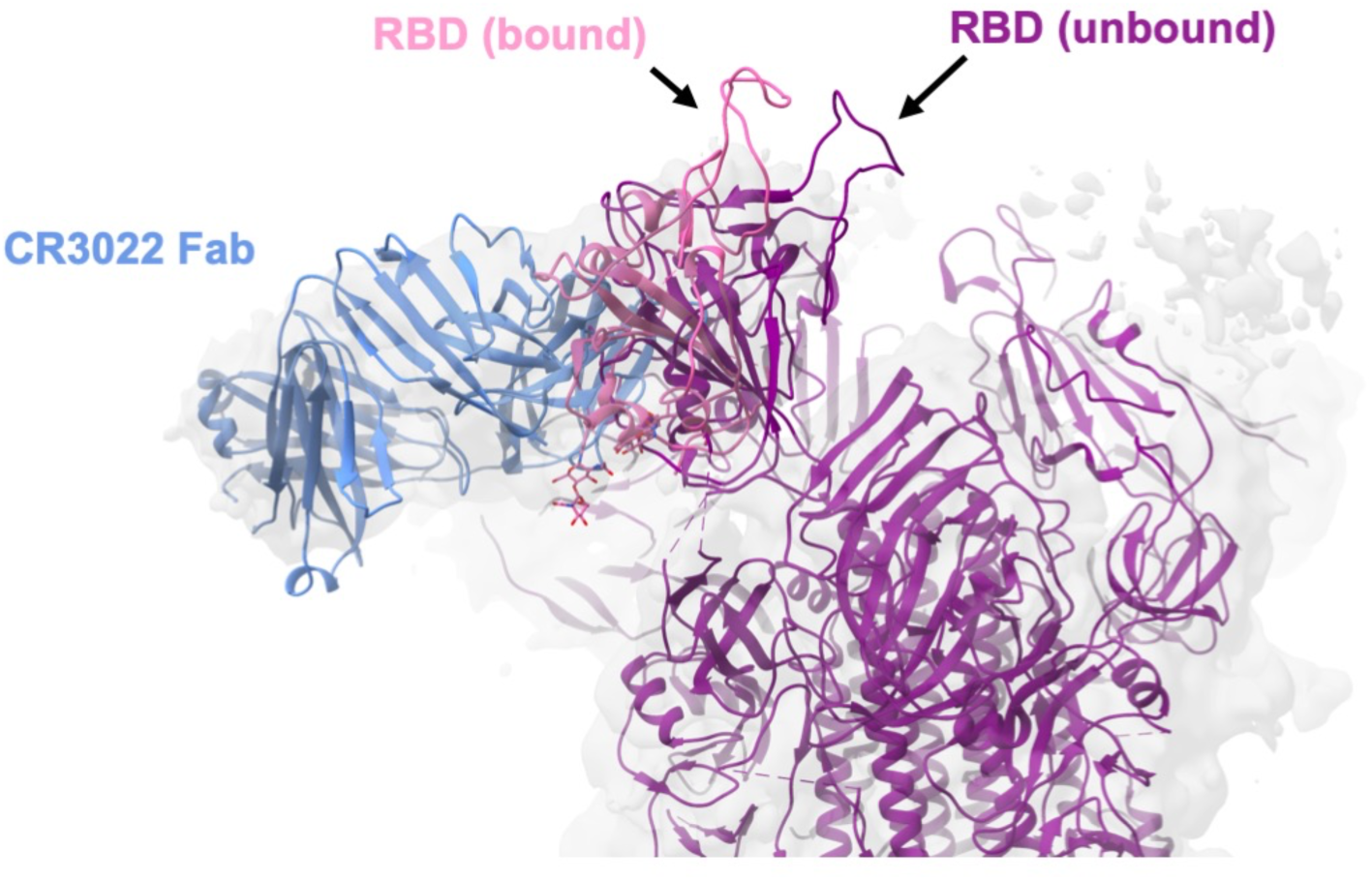
Comparison of conformations of CR3022-bound and unbound RBDs. The conformation of CR3022-bound RBD in class 2 and 4 is compared to the conformation of RBD on an unliganded SARS-CoV S protein (PDB 6ACD) [35].

**Supplementary Figure 6.**
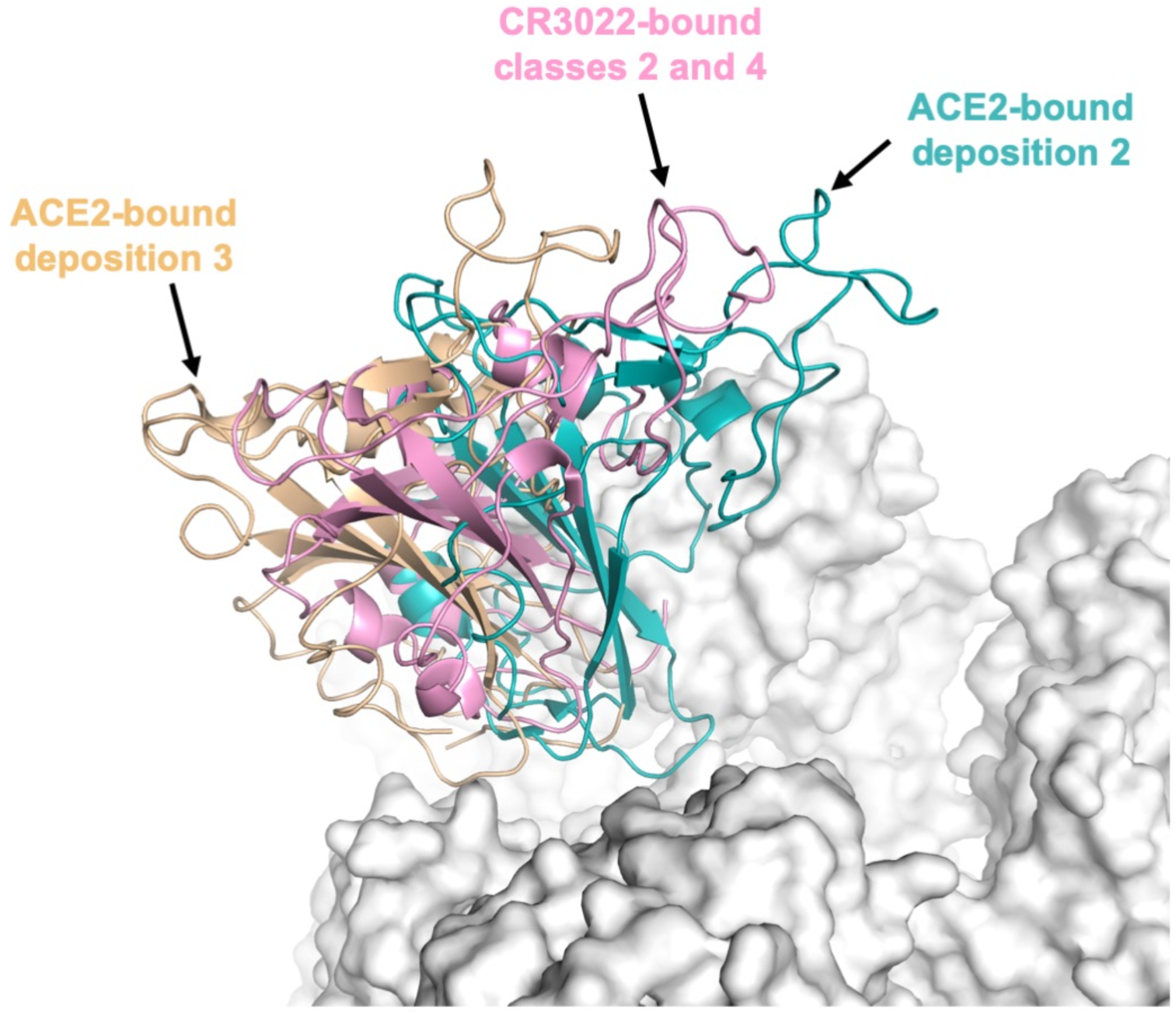
Comparison of conformations of CR3022-bound and ACE2-bound RBDs. The conformation of CR3022-bound RBD in class 2 and 4 is compared to that of depositions 2 and 3 of ACE2-bound RBD (PDB 6ACJ and 6ACK, respectively) [35].

